# Harnessing AI to Build Virtual Cells

**DOI:** 10.64898/2026.04.11.717183

**Authors:** Xingyi Cheng, Pan Li, Han Guo, Youwei Liang, Jing Gong, William de Vazelhes, Changjiang Gou, Pengtao Xie, Le Song, Eric Xing

## Abstract

A virtual cell is a world model of a cell: a computational system that predicts, simulates and programs cellular processes across modalities and scales. An important path toward this goal is to model how genetic and chemical perturbations give rise to transcriptional responses, a core capability for disease understanding and drug discovery. However, current approaches remain expert-intensive, relying on iterative manual model design, training and debugging over months.

Here we present VCHarness, an autonomous AI system that constructs perturbation-response models by combining an AI coding agent with multimodal biological foundation models. The system explores large spaces of architectures and training pipelines with minimal human intervention, iteratively generating, evaluating and refining candidate models. Across multiple perturbation-response benchmarks, VCHarness identifies architectures that outperform expert-designed approaches while reducing development time from months to days. It further uncovers non-obvious architectural patterns associated with improved performance, indicating that automated search can extend beyond conventional design strategies. These results suggest a shift from manually engineered models toward autonomous systems for constructing components of virtual cell world models, enabling scalable and data-driven exploration of cellular systems.

## 1 Introduction

A virtual cell is a world model of a cell: a computational system that predicts, simulates and programs multimodal cellular processes across scales by integrating heterogeneous and multi-modal data within a unified framework [1]. Such models will enable systematic exploration of cellular function and intervention, with direct implications for disease modeling, target identification and drug discovery [2, 3, 1].

A step toward this challenging goal begins with more limited but important capabilities, such as predicting gene expression changes under genetic or chemical perturbations [4, 5, 6, 7, 8, 9]. Although this does not capture the full scope of a virtual cell, it represents a core functionality that is already useful in practice. Accurate models of perturbation-induced transcriptional change therefore serve as a foundational building block for more general virtual cell systems.

In practice, however, even this relatively constrained task remains slow and expert-intensive. Model development still depends on hand-designed architectures, repeated debugging, manual hyperparameter tuning and long cycles of empirical iteration. Benchmark studies further indicate that performance is highly sensitive to design choices that are difficult to enumerate exhaustively [10]. As a result, progress is constrained not only by data and compute, but also by the limited bandwidth of human-guided search over model designs.

At the same time, a convergence of technical trends points toward a different paradigm. Foundation models provide reusable building blocks for biological representation learning and strong initialization that can reduce the effective search burden [11, 12, 5, 6, 7]. Coding agents can synthesize, modify and debug machine learning pipelines from natural-language task descriptions [13, 14, 15, 16]. Structured search algorithms such as Monte Carlo tree search (MCTS) offer principled mechanisms for allocating exploration across large decision spaces [17]. At a systems level, recent works on automated program search and AI harness systems suggest that closed-loop agentic frameworks can iteratively construct solutions that extend beyond conventional human design heuristics [18, 19, 20, 21].

These developments suggest that the construction of virtual cell components, such as perturbation-response models, can itself be automated, transforming model building from a manual, expert-driven process into a data-driven search over executable programs. Accelerating the development of reliable perturbation-response models is therefore not only a practical objective, but also a necessary step toward more general, world-model–like representations of cellular systems.

Here we present VCHarness, an autonomous AI system for perturbation-response model construction. VCHarness combines multimodal biological foundation-model components, an AI coding agent, Monte Carlo tree search, structured feedback memory and distributed execution into a closed-loop system over executable programs. Rather than optimizing within a fixed architecture family, VCHarness searches over complete training pipelines, including data processing, multimodal foundation models, architectural composition and optimization strategies. We demonstrate across multiple CRISPR knockdown-response tasks that VCHarness outperforms strong expert-designed baselines, operates with measurable efficiency, and discovers non-obvious architectural patterns supported by search analysis and ablation studies.

Taken together, this work suggests a path toward accelerated construction of virtual cell systems that not only predict biological responses but also support the construction of increasingly comprehensive and coherent world models of cellular behavior. VCHarness may enable scalable generation of virtual cells across diverse biological contexts, with potential implications for accelerating discovery, guiding experimental design and rethinking the role of human expertise in computational biology.

## 2 System Overview

VCHarness is an end-to-end autonomous system for perturbation-response model construction. Rather than fixing an architecture and tuning its parameters, it treats the entire model-building workflow— program generation, debugging, execution, evaluation and refinement—as a single searchable loop (Fig. 1). For each task, the system takes as input the dataset, the task objective and lightweight natural-language specifications, and iteratively produces candidate programs without manual intervention in model design.

**Fig. 1.**
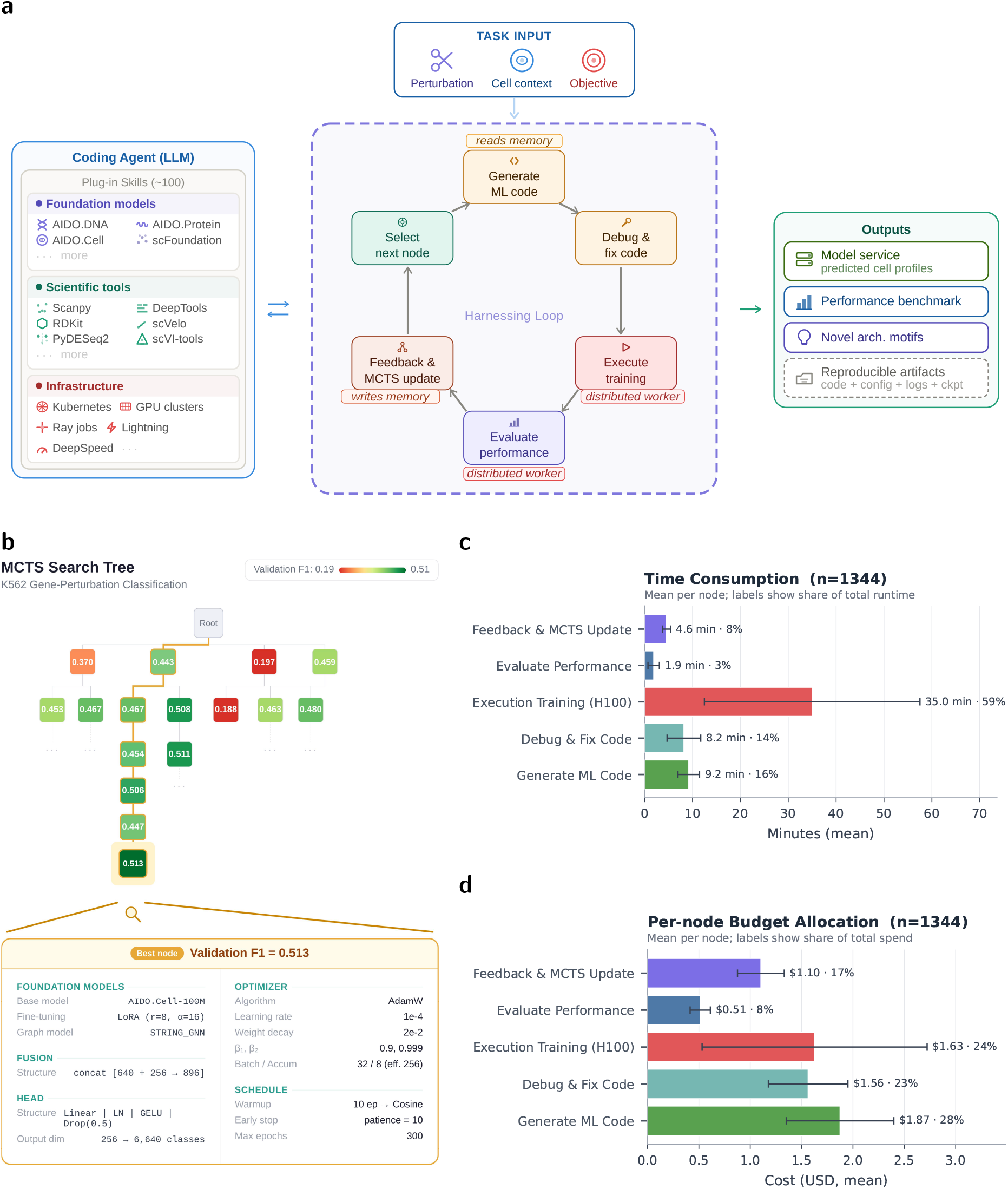
Overview of the VCHarness system. **a**, The system integrates the AIDO family of biological foundation models, a coding agent, roughly 100 plug-in skills for virtual-cell model development, and Monte Carlo tree search (MCTS)-based node selection. It is a distributed system that can train multiple models in parallel while orchestrating the overall search process across workers. **b**, Representative MCTS search tree for K562 differential gene expression prediction task. Node color indicates validation F1 score, and the highlighted trajectory shows iterative refinement toward higher-performing regions of program space. The lower panel summarizes the best-performing configuration (validation F1 = 0.513), including used foundation models, fusion strategy, prediction head and optimization settings. **c**, Mean stage-wise runtime per MCTS node, aggregated across K562 search runs. Model execution and training dominate runtime, whereas evaluation, feedback and tree updates contribute comparatively little overhead. **d**, Mean stage-wise budget allocation per MCTS node (USD), aggregated across K562 search runs. Execution training (H100) is estimated using GPU-hour pricing, while other stages follow Claude Sonnet 4.6 pricing. Budget is distributed across program generation, debugging, and execution training at comparable scale, whereas evaluation contributes a smaller share.

At its core, VCHarness combines the AIDO family of biological foundation models, an AI coding agent and Monte Carlo tree search (MCTS) into a unified system. The AIDO models provide generalizable representations and modules for biological data, while MCTS organizes the search space over executable programs, balancing exploration of new model designs with refinement of promising candidates. The coding agent generates candidate programs, composes AIDO-based components into architectures, debugs execution failures, and summarizes prior programs and training outcomes to guide subsequent proposals. These steps are integrated within a single harness in which generated programs are executed, evaluated and inserted into a shared search tree that conditions future exploration (Fig. 1a,b). Each generated architecture is assembled with a consistent three-stage layout—foundation-model encoders, a fusion module that blends heterogeneous signals, and a task-specific prediction head—so that the search can focus on which foundation-model/fusion/head combinations unlock new generalization behavior rather than juggling entirely bespoke layouts from scratch.

At the system level, this design enables continuous refinement over a large space of model architectures and training pipelines. In practice, the coding agent is augmented with roughly 100 built-in skills specialized for virtual-cell modeling, including reusable tools for biological foundation-model integration, data preprocessing, distributed training launch, evaluation, debugging, and result summarization, as illustrated in Fig. 1a. At the per-node level in the search tree, model training and execution dominate runtime and cost, while evaluation, feedback and tree updates introduce comparatively modest overhead (Fig. 1c,d). This concentration of compute on model training improves overall efficiency.

In summary, VCHarness is an operationally tractable system capable of supporting day-scale autonomous exploration over large model spaces.

## 3 Results

**VCHarness achieves consistent performance gains across diverse datasets** We first asked whether VCHarness can achieve consistent improvements compared with human experts across heterogeneous perturbation-response datasets [22, 23]. As illustrated in Fig. 2, VCHarness consistently outperforms all expert-designed baselines across CRISPR knockdown DEG-classification tasks in four distinct cell lines (HepG2, Jurkat, K562, and hTERT-RPE1). Specifically, VCHarness achieves a mean Macro-F1 ranging from 0.4423 to 0.4761, marking a significant improvement over the GNN Simple baseline, which fluctuates between 0.3966 and 0.4470 (Fig. 2a).

**Fig. 2.**
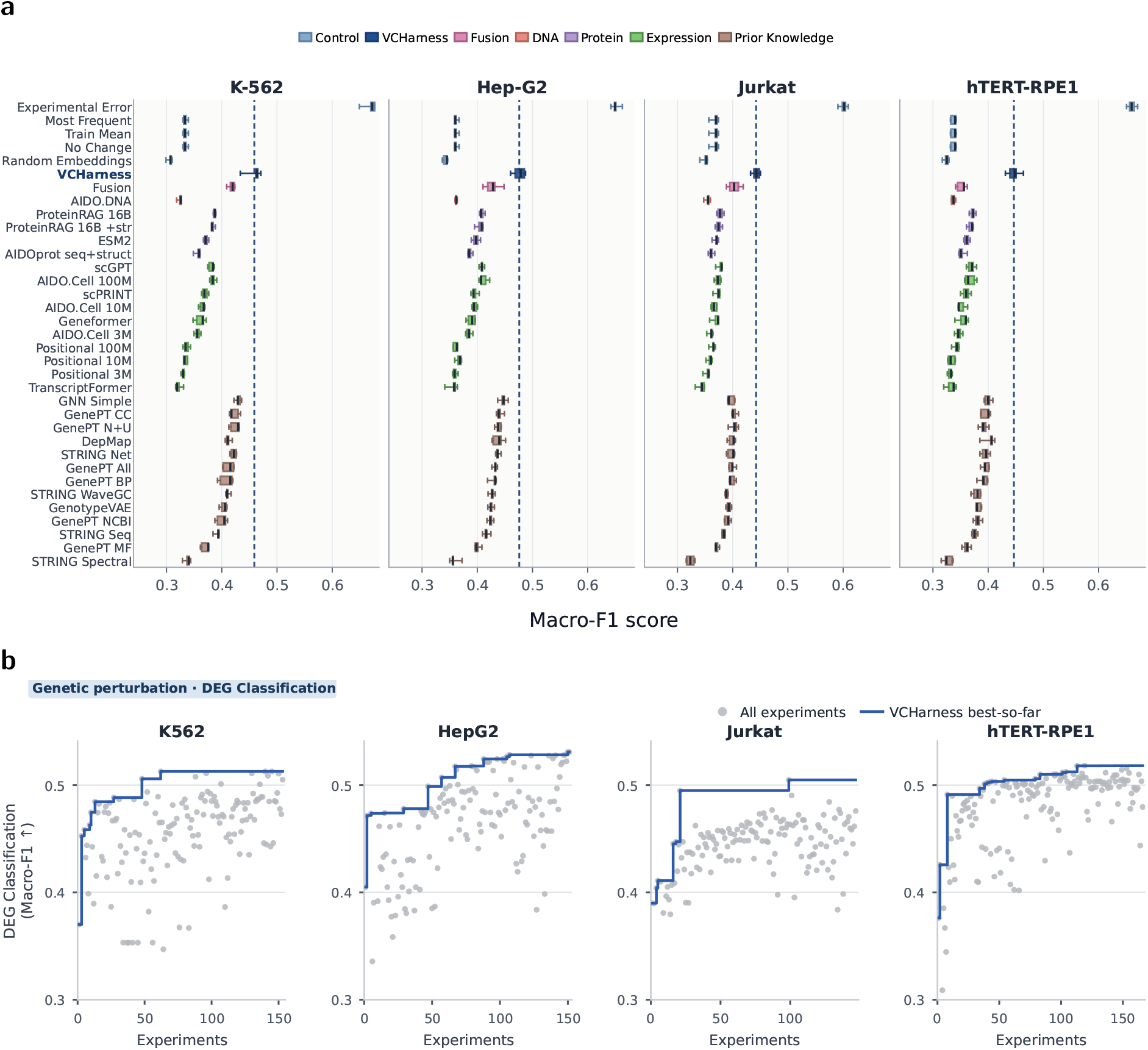
VCHarness performance and search dynamics across CRISPR perturbation settings. **a**, VCHarness (blue) outperforms human-designed models grouped by input modality or knowledge source across four differentially expressed gene (DEG) classification tasks on four cell lines (HepG2, K562, Jurkat, and hTERT-RPE1); dashed lines indicate mean performance. Baselines are from the foundation-models-perturbation repository (https://github.com/genbio-ai/foundation-models-perturbation/tree/main/results/scores). Consistent gains across cell lines demonstrate transferability of the autonomous search loop across cell lines. The *y*-axis corresponds to test macro-F1 scores. **b**, Best-so-far trajectories during Monte Carlo tree search (MCTS). Grey dots denote evaluated candidates, and blue lines indicate the running best. Rapid early improvement followed by gradual refinement reflects efficient allocation of search effort. The *x*-axis represents candidate implementations ordered by evaluation time, and the *y*-axis denotes validation macro-F1 scores.

This result is notable because these tasks differ substantially in cell type, perturbation context and data characteristics. Consistent improvements across all settings suggest that the gains do not arise from a single architecture tuned to a specific benchmark, but from a more generalizable model-construction process. The same autonomous search loop can be applied across datasets while still discovering task-appropriate solutions, supporting the view that VCHarness provides a transferable system for searching over large model design spaces rather than producing fixed architectures tuned to a single benchmark.

### VCHarness identifies highly performant models efficiently

To characterize the optimization dynamics of VCHarness, we examined the best-so-far performance trajectories (Fig. 2b). Across all tested cell lines, the framework demonstrates a consistent upward trajectory, with rapid initial gains followed by steady refinement. Specifically, while the best seed implementation for each cell line initially achieves a Macro-F1 of approximately 0.440, the iterative search within the harnessing loop successfully elevates performance to roughly 0.510. This measurable improvement underscores the critical role of the harnessing loop in iteratively refining and optimizing task-specific implementations beyond their initial baselines.

This behavior is consistent with the expected dynamics of Upper Confidence Bound (UCB)-guided Monte Carlo tree search (MCTS) [17]. Initial exploration promotes broadly sampling the program space to identify promising regions, after which computation is progressively concentrated on trajectories that continue to yield improvements. The resulting trajectories suggest that VCHarness derives efficiency from structured allocation of search effort, rather than from random trial-and-error over disconnected model variants.

### VCHarness discovers novel and non-obvious model architectures

Aggregate benchmark scores do not fully capture what VCHarness discovers, so we next examined the resulting architectures. Fig. 3 presents a case study on the hTERT-RPE1 differential gene expression prediction task, where VCHarness identifies a significantly more accurate architecture that lifts validation Macro-F1 from 0.3445 to 0.5182. This discovered model combines graph neural networks [10] over the STRING database [24] with selective fine-tuning and perturbation-conditioned computation, moving beyond simpler, modality-isolated templates.

**Fig. 3.**
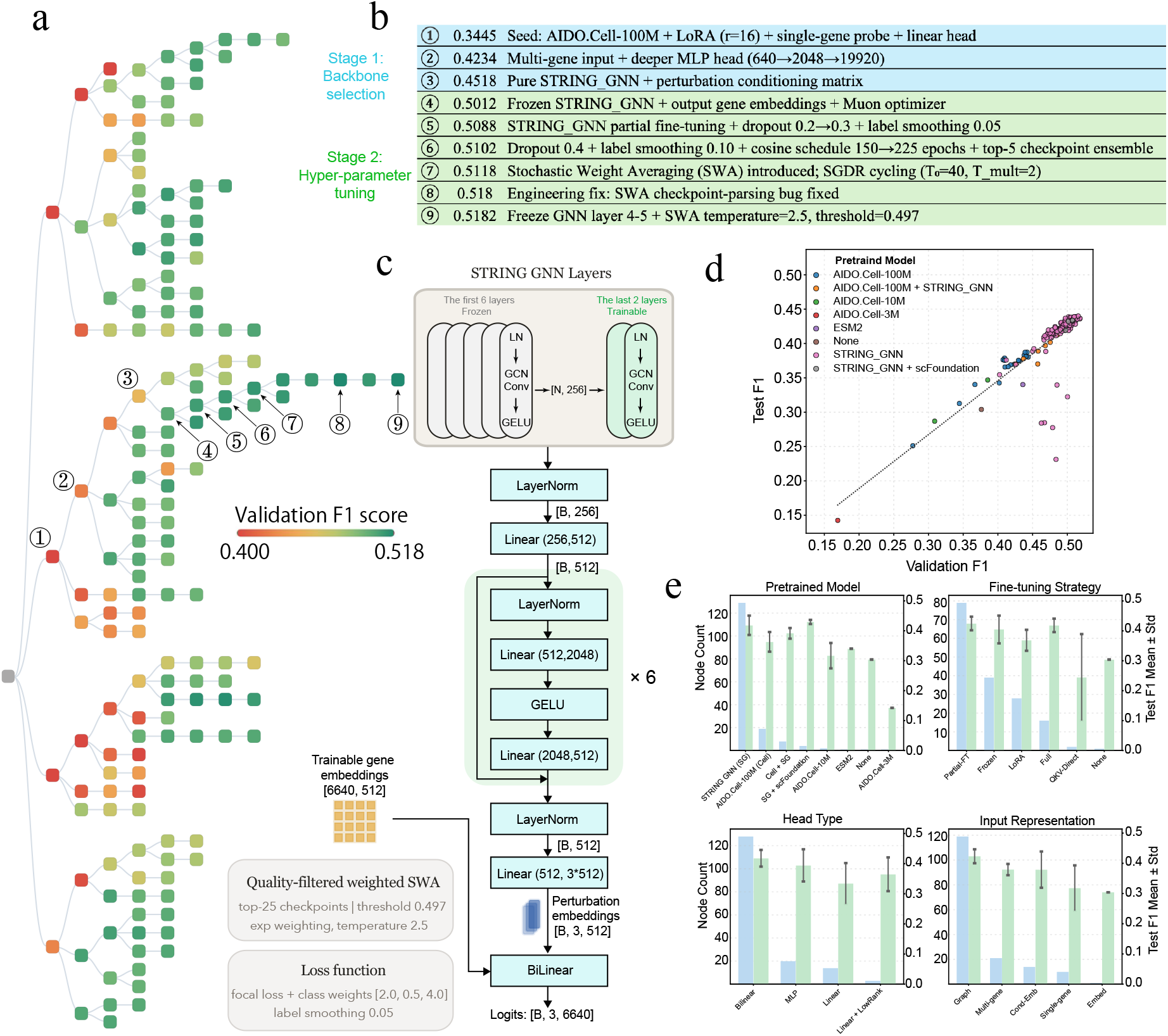
VCHarness discovers non-obvious architectural patterns through search. **a**, MCTS visualization for the hTERT-RPE1 DEG-classification campaign. Node color encodes validation macro-F1, and the highlighted trajectory shows how search progressively concentrates on higher-performing regions of program space. **b**, Representative local edits made along the successful branch in **a**. These ranked modifications illustrate how performance gains arise from sequential architecture and optimization changes rather than from a single one-step jump. **c**, Architecture recovered at the node marked ⑨ in **a**. The discovered model combines graph structure, selective fine-tuning, and perturbation-conditioned computation, illustrating how VCHarness assembles executable programs that differ from simpler modality-isolated baselines. **d**, Correlation between validation and test scores across evaluated nodes. The near-linear agreement indicates that validation-guided selection generalizes reliably to held-out test data. **e**, Aggregate motif analysis over searched architectures. By grouping discovered models according to frequently occurring design choices, this panel highlights neural-architecture motifs that are repeatedly associated with stronger performance.

As the MCTS visualization tree (Fig. 3a) illustrates, the framework iteratively improves upon the initial seeds through systematic exploration. Specifically, the most promising lineage is identified early (Stage 1) and is subsequently optimized by exploring architectural variants and hyper-parameter configurations that branch from these high-performing parent nodes (Stage 2, Fig. 3b). Additional cell-line-specific discovery examples for HepG2, K562, and Jurkat are provided in the Appendix.

This analysis highlights a key qualitative result: VCHarness does not simply retune standard expert baselines. Instead, it emerges a stage-wise search strategy to assemble novel and non-obvious combinations of components whose contributions are supported by ablation and by local search-tree structure (Fig. 3b, c). In the hTERT-RPE1 example, partial fine-tuning [25] outperforms full fine-tuning, suggesting that constraining updates to selected graph layers can preserve useful pretrained structure while avoiding excessive smoothing [26]. A detailed description of the model architecture corresponding to Fig. 3c is provided in Algorithm 1.

Across cell lines, the discovered models exhibit recurring patterns, including hybrid encoders, multi-branch feature extraction and context-dependent fusion. The positive correlation between validation and test performance across searched nodes further supports the use of validation score as a practical search objective during autonomous exploration (Fig. 3d). In addition, motif-frequency analysis over the searched architectures reveals repeatedly recovered neural-architecture patterns rather than isolated one-off designs (Fig. 3e). Together, these observations indicate that agent-based search can uncover effective architectural strategies that are difficult to identify through manual workflows alone.

## 4 Methodology

### Problem formulation

VCHarness treats perturbation-response model construction as a search problem over executable machine learning pipelines. Given a perturbation-response dataset 𝒟 and an evaluation metric *J*(·), the goal is to identify a model *f*_*θ*_ with strong empirical performance while operating under a finite compute budget. Rather than searching over parameters within a fixed architecture family, VCHarness searches over programs that jointly specify data processing, representation modules, architectural composition, training configuration, and optimization strategy. This formulation makes model development itself the object of search and aligns the optimization target with the actual practice of model building, where architectural edits, debugging decisions, and training choices all affect final performance.

### Closed-loop model construction

The system operates as an iterative closed loop that alternates between proposal, debugging, execution, evaluation, and refinement. At iteration *t*, VCHarness selects a candidate program from the current search state according to the MCTS UCB score and uses the coding agent to generate a modified program while reading search history from prior branches and evaluations. The resulting program is debugged and then executed to obtain a trained model and evaluation signal, after which the system writes run feedback back into the shared memory and updates the search graph with the new outcome. Subsequent proposals are therefore conditioned not only on the current branch, but also on accumulated search history and run feedback from earlier evaluations. In this way, empirical results directly shape future model construction decisions, following the broader pattern of agentic systems that interleave generation, execution, and feedback rather than relying on one-shot synthesis [27, 20].

### Program representation and search space

Each node in the search process corresponds to a program *P* representing a complete machine learning pipeline. A program may include data preprocessing steps, choices of biological foundation-model encoders, fusion operations, task heads, loss functions, and training schedules. Child programs are generated by modifying parent programs, which allows VCHarness to build increasingly sophisticated candidates through incremental edits rather than repeated construction from scratch.

The search structure is primarily tree-based, but in practice supports **DAG-like** reuse of successful ancestors and shared design motifs. This representation is useful because many high-performing candidates are not independent inventions; they are compositional refinements of earlier programs that already encode partially successful design choices.

### Biological foundation-model library

The search space is anchored by a reusable library of pretrained biological components spanning multiple representational levels. In the current system, these include inhouse genome, RNA, protein, interactome and cell foundation-model variants such as AIDO.DNA [28], AIDO.DNA2 [29], together with AIDO.Protein [30] and AIDO.Cell [31], as well as public protein and single-cell models such as ESM2 [11], Geneformer [5], scGPT [7], and scFoundation [6], and graph-based components such as STRING GNN [24]. VCHarness treats these components as modular primitives that can be selected, composed, partially fine-tuned, or combined with task-specific heads and fusion layers. This design follows recent work showing that large-scale biological pretraining can provide broadly useful priors for downstream cellular tasks [28, 11, 5].

Formally, each pretrained component can be viewed as a mapping *φ*_*m*_ : 𝒳_*m*_ → ℝ^*d*^, where 𝒳_*m*_ is the modality-specific input domain and ℝ^*d*^ is a learned representation space. The role of VCHarness is not simply to choose one backbone from this library, but to search over how such components should be adapted and fused inside executable programs.

### Monte Carlo tree search

The process of finding a high-performing model is organized in a tree data structure. To allocate search budget efficiently, VCHarness uses Monte Carlo tree search (MCTS) to balance exploration and exploitation over program space. Model discovery is therefore organized not as a flat queue of independent trials, but as a structured sequence of expansions from previously evaluated programs.

At each decision point, VCHarness selects nodes using an upper confidence bound criterion,

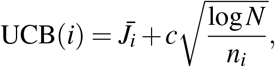

where 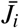 is the average score of node *i, n*_*i*_ is its visit count, *N* is the visit count of its parent node, and *c* controls the exploration–exploitation trade-off. Selected nodes are expanded through coding-agent edits, executed, and their outcomes are propagated back into the search state.

This coupling between MCTS and code generation is central to VCHarness: the search policy determines where to explore, while the coding agent determines which concrete program to instantiate in that region. At any iteration *t*, the set of previously evaluated programs {*P*_*i*_*}*_*i<t*_ and their corresponding scores {*J*_*i*_*}*_*i<t*_ define the search history ℋ_*t*_, which conditions subsequent exploration.

### AI coding agent

The coding agent parameterizes a policy *π*(*P* |ℋ_*t*_) over candidate programs conditioned on the search history ℋ_*t*_ = {(*P*_*i*_, *J*_*i*_)}_*i<t*_. Given the task description, prior code, and aggregated feedback from previous evaluations, the agent produces executable code specifying model architecture, training configuration, and data handling logic.

In practice, the agent serves four roles. First, it performs program synthesis by generating new candidate programs. Second, it performs execution repair by diagnosing and correcting runtime or logical failures. Third, it summarizes evaluation outcomes into structured memory records that capture successful motifs, failure modes, and useful implementation details. In our implementation, these records are written after each run as run feedback and then aggregated across the current search tree before the next expansion step, so proposals can read search history from the full tree rather than only from a single ancestral path. Fourth, it performs iterative refinement by incorporating those records into subsequent proposals. This design allows the search process to accumulate reusable experience instead of treating each candidate as an isolated sample. We rely on Claude Sonnet 4.6 as the coding agent that performs these roles, both for program generation and for the structured feedback synthesis described above.

### Evaluation and feedback

Each candidate program *P*_*t*_ is executed to produce a trained model 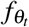 together with an evaluation score *J*_*t*_. Depending on the task, the recorded signals may include predictive metrics such as micro- or macro-F1 or task-specific loss, as well as runtime, failure traces, and monetary cost. These signals serve two purposes. They update the search statistics used by MCTS, and they supply the feedback memory consumed by the coding agent when generating future proposals. Evaluation is therefore not only a terminal judgment step, but also a mechanism for shaping subsequent search behavior. This makes the loop empirically grounded: future proposals are influenced by what actually ran successfully, what failed, and what yielded useful gains under the available compute budget.

### Distributed execution

Because candidate model execution is the dominant cost in the loop, VCHarness is designed to run in a distributed environment across multiple workers and GPUs. Worker nodes asynchronously execute selected programs, while a shared state maintains the global search structure, node statistics, and experiment history. Synchronization ensures consistency across parallel updates, and job isolation allows individual failures to be handled locally without collapsing the larger campaign. At a systems level, this execution model resembles recent harness-based agent frameworks in which generation, tool use, and validation are coordinated through a persistent control loop rather than through isolated script calls [21, 32, 33]. This infrastructure is essential for turning VCHarness from a toy search procedure into a system capable of evaluating large numbers of candidate programs robustly under realistic time constraints.

### Summary

Taken together, these components define VCHarness as a robust and efficient system for autonomous cellular perturbation-response model construction. Biological foundation models provide reusable representation primitives, the coding agent converts search decisions into executable programs, MCTS allocates exploration budget, evaluation produces both scores and actionable feedback, and distributed execution makes the overall process tractable. The result is a system in which empirical performance continually drives iterative program refinement.

## 5 Discussion

VCHarness demonstrates that cellular perturbation response models can be constructed through an autonomous system rather than exclusively through manual expert iterations. The results suggest that strong performance emerges not from a single handcrafted architecture, but from the interaction of reusable biological building blocks, coding-agent-driven program synthesis and other guardrails such as debugging, structured search, and iterative feedback. In that sense, VCHarness is closer to emerging agentic scientific systems and autonomous discovery pipelines than conventional AutoML over a fixed hypothesis class [20, 34, 35, 36]. At the same time, the current system exposes several aspects to improve that define better future systems.

### Experience reuse across datasets and problems

VCHarness accumulates substantial empirical history during a campaign, but this history is still only partially reusable across tasks. Feedback summaries capture useful local information, yet they do not fully support transfer of architectural motifs, debugging strategies, or successful optimization patterns across domains. Notably, STRING GNN-related designs consistently appear among top-performing nodes across all four cell lines (Fig. 3d, 4d, 5d, 6d), suggesting that shared architectural motifs do emerge and could be explicitly leveraged for cross-task transfer. Layered and structured memory representations, reusable code abstractions, and more explicit cross-task retrieval mechanisms may further improve sample efficiency and reduce redundant exploration in future campaigns.

### Biological prior knowledge can be used for better guidance or constraints

The current methodology is primarily data-driven. This makes it flexible and often effective, but it can also produce architectures whose biological interpretation is less direct than their predictive performance [37]. Interestingly, the most performant strategies consistently opted for GNN architectures leveraging STRING PPI data (Fig. 3d), suggesting that even a data-driven search can recover biologically grounded inductive biases. Incorporating more comprehensive curated pathway knowledge, mechanistic constraints, structural priors from AlphaFold-class systems, or literature-grounded proposals—each of which could be implemented as a reusable skill—could further guide search toward solutions that are not only accurate but also more interpretable and biologically plausible [38, 39].

### The search policy can be further improved

MCTS provides a strong and general-purpose mechanism for balancing exploration and exploitation, but it is unlikely to be the final word on search over biological program space. More structured proposal distributions, learned edit policies, hierarchical search spaces, or multi-fidelity evaluation strategies may further improve efficiency. A particularly natural extension would be explicitly hierarchical search that first allocates budget over coarse architectural decisions—encoder family, fusion pattern, or head design—and only then refines training hyperparameters within the most promising structural families. This differs from the stage-wise refinement already visible in Fig. 3b, which emerges organically from MCTS rather than being enforced by the search policy; a hierarchical policy would impose this structure explicitly and more efficiently. In particular, tighter coupling between search policy and historical experience could allow VCHarness to avoid repeatedly exploring low-value regions of program space, similar to how recent agentic ML systems increasingly combine exploration with explicit reasoning and longer-horizon planning [40, 41, 33].

### Compute cost can be further optimized

Although per-node measurements indicate that VCHarness is operationally tractable, execution still dominates both runtime and spend. This is a practical reminder that autonomous model construction remains constrained by evaluation cost. On the model-evaluation side, techniques such as early stopping, surrogate scoring, partial training, staged filtering of candidates, or tighter integration with self-driving-lab style resource schedulers could reduce cost while preserving the benefits of structured search. On the agent side, additional savings may come from heterogeneous model allocation across stages of the loop: expensive frontier models may be reserved for proposal generation, difficult debugging, or major architectural revisions, whereas lower-cost or open-source models may be sufficient for feedback summarization, run-log parsing, memory consolidation, or other routine bookkeeping steps.

### Broader validation for system effectiveness is needed

The present study focuses on perturbation-response prediction, particularly CRISPR-based settings. This is an important domain, but it is only one slice of the broader virtual cell problem. Extending VCHarness to temporal dynamics, multimodal response prediction, cell–cell interactions, or multiscale cellular behavior will require richer datasets, stronger open benchmarks, and more expressive search spaces [42].

Overall, these limitations do not undercut the central result. Instead, they provide clear routes and methodology for VCHarness for further improvement and development as a scalable framework for constructing virtual cells. The practical challenge now is not whether such systems can work at all, but how to make them more reusable and robust, better grounded in biology, and more rigorous under realistic evaluation constraints.

## 6 Conclusion

We presented VCHarness, an autonomous AI system for cellular perturbation-response model construction that harnesses biological foundation-model components, coding-agent-based program synthesis, Monte Carlo tree search, feedback memory, and distributed execution. By framing model development as search over executable programs, VCHarness moves the optimization target from a fixed architecture class to the broader space of trainable programs.

Across multiple perturbation-response tasks, VCHarness discovers models that outperform strong expert-designed baselines while operating within day-scale autonomous campaigns. The system also exhibits measurable per-node efficiency and uncovers non-obvious architectural patterns that are supported by search trajectories and ablation. These findings suggest that autonomous model construction can become a practical methodology for scalable virtual cell research rather than merely a proof-of-concept automation tool.

Looking ahead, VCHarness points toward a shift in computational biology workflows: experimental data and evaluation objectives remain central, but the search over model designs can increasingly be delegated to structured autonomous systems. Relative to recent efforts in automated scientific research and agentic biological modeling [18, 21, 40, 41], VCHarness emphasizes executable model construction under biological evaluation objectives as the primary object of search. Extending this paradigm to richer biological settings, stronger priors, and more efficient search policies is a natural next step toward more general autonomous scientific discovery systems.

## A Supplementary analyses

The main text establishes that VCHarness improves over strong baselines across multiple CRISPR perturbation-response settings. This Appendix extends that result by providing additional architecture-discovery case studies for the remaining cell lines from the Essential dataset [23, 22, 10], supplementing the hTERT-RPE1 example shown in the main text (Fig. 3).

### A.1 Discovered Architecture Across Essential Cell Lines

The diversity of optimal architectures discovered by VCHarness across different cell lines underscores the necessity of cell-specific model tailoring. While the main text details the hTERT-RPE1 case (Fig. 3, Alg. 1), analogous search campaigns for HepG2, Jurkat, and K562 reveal distinct evolutionary paths and architectural motifs (Figs. 4–6, Algs. 2, 3, and 4).

**Fig. 4.**
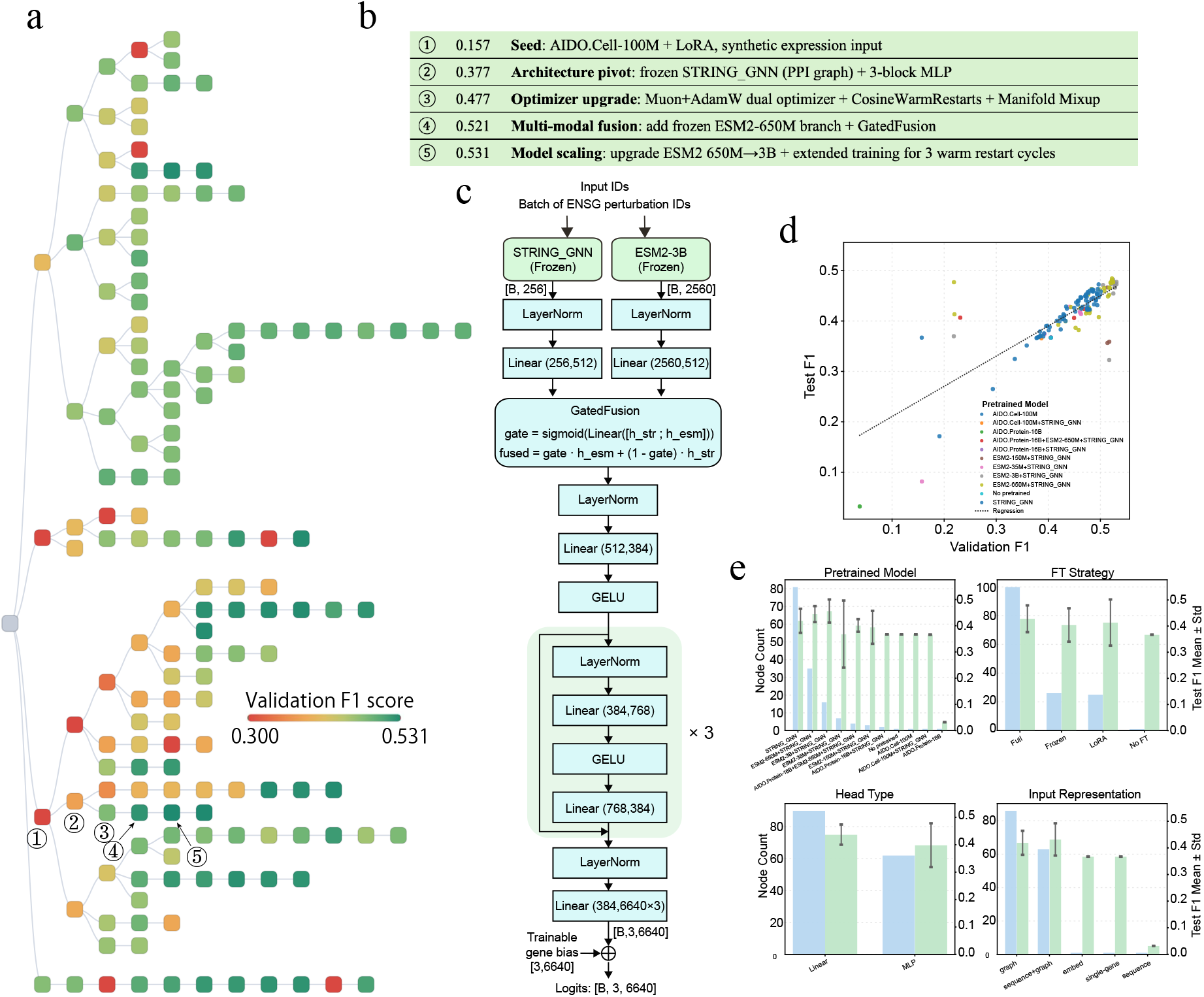
Architecture discovery on HepG2. **a**, MCTS visualization for the HepG2 Essential campaign. **b**, Ranked local modifications along a successful branch, showing how search improves performance through sequential edits. **c**, Best-discovered model architecture recovered from the highlighted branch. **d**, Correlation between validation and test scores across evaluated nodes, supporting the use of validation performance as a practical search objective. **e**, Aggregate motif analysis over searched architectures, summarizing which neural-architecture choices recur most often among stronger HepG2 models. In this campaign, the best-performing solutions repeatedly favored multimodal fusion centered on graph-based perturbation structure with additional pretrained expression features.

**Fig. 5.**
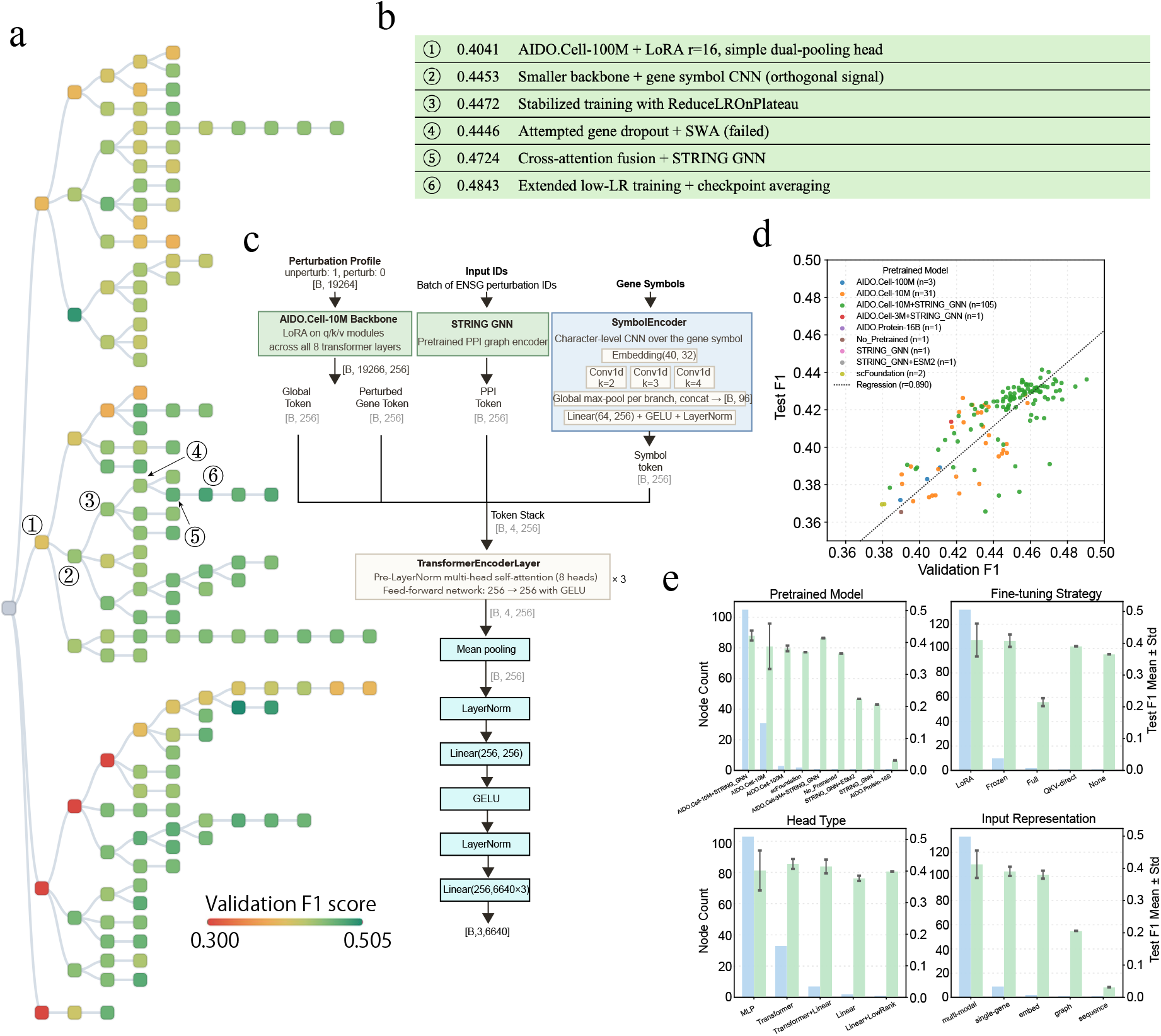
Architecture discovery on Jurkat. **a**, MCTS visualization for the Jurkat Essential campaign. **b**, Ranked local modifications along a successful branch. **c**, Best-discovered Jurkat model architecture. **d**, Correlation between validation and test scores across evaluated nodes. **e**, Aggregate motif analysis over searched architectures, highlighting recurrent neural-architecture motifs associated with stronger Jurkat performance. Relative to HepG2, the stronger Jurkat nodes more often relied on compact architectures with explicit perturbation modeling and lighter fusion strategies.

**Fig. 6.**
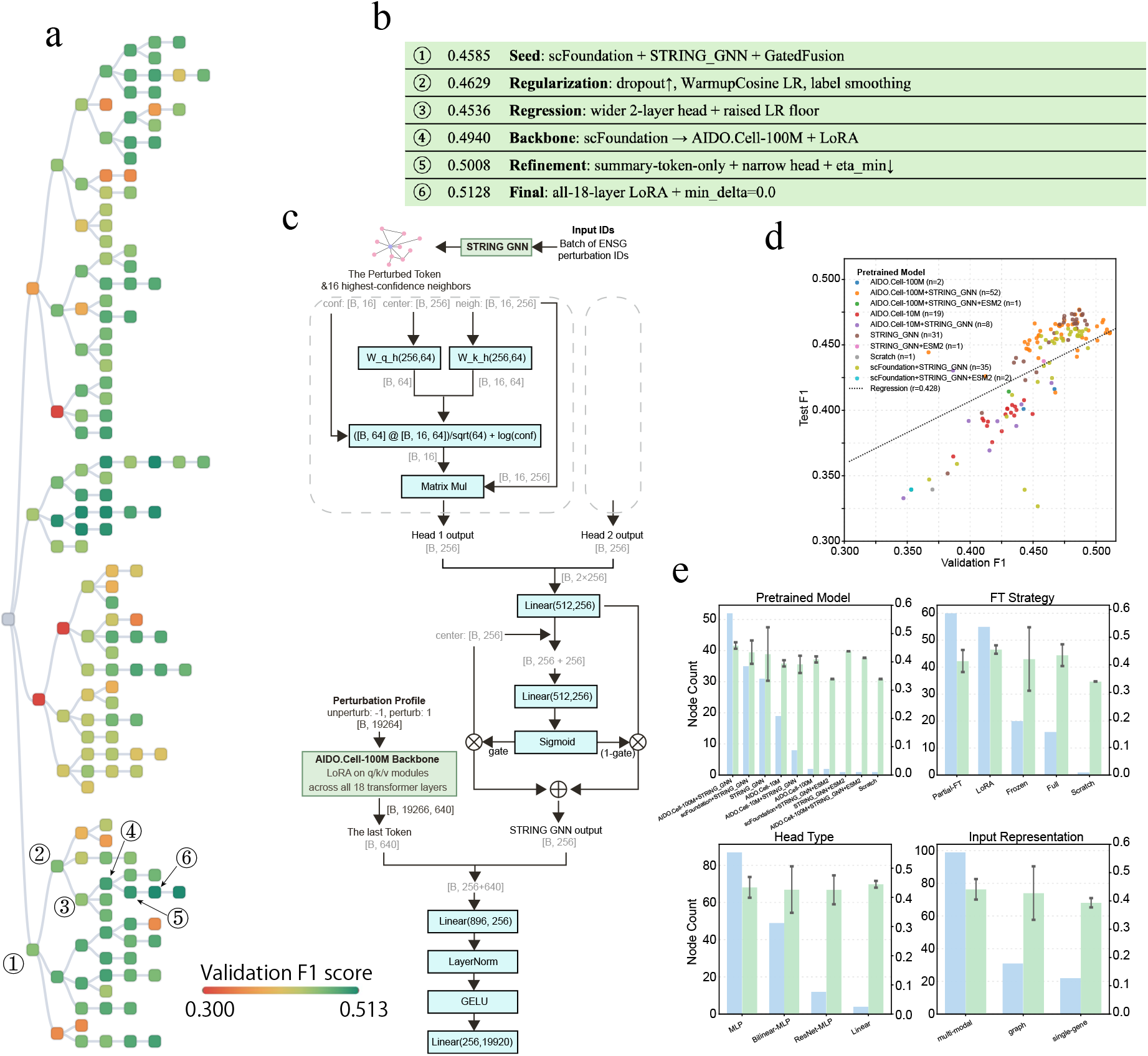
Architecture discovery on K562. **a**, MCTS visualization for the K562 Essential campaign. **b**, Ranked local modifications along a successful branch. **c**, Best-discovered K562 model architecture. **d**, Correlation between validation and test scores across evaluated nodes. **e**, Aggregate motif analysis over searched architectures, summarizing recurring architectural patterns associated with improved K562 performance. In K562, successful nodes repeatedly combined strong cell-level backbones with graph-based perturbation modules and gated fusion, suggesting a particularly stable interaction between pretrained cellular representations and explicit network priors.

#### Algorithm 1

Discovered Architecture — hTERT-RPE1 Perturbation DEG (Reverted Partial STRING-GNN + Deep Bilinear MLP + Quality-Filtered SWA)

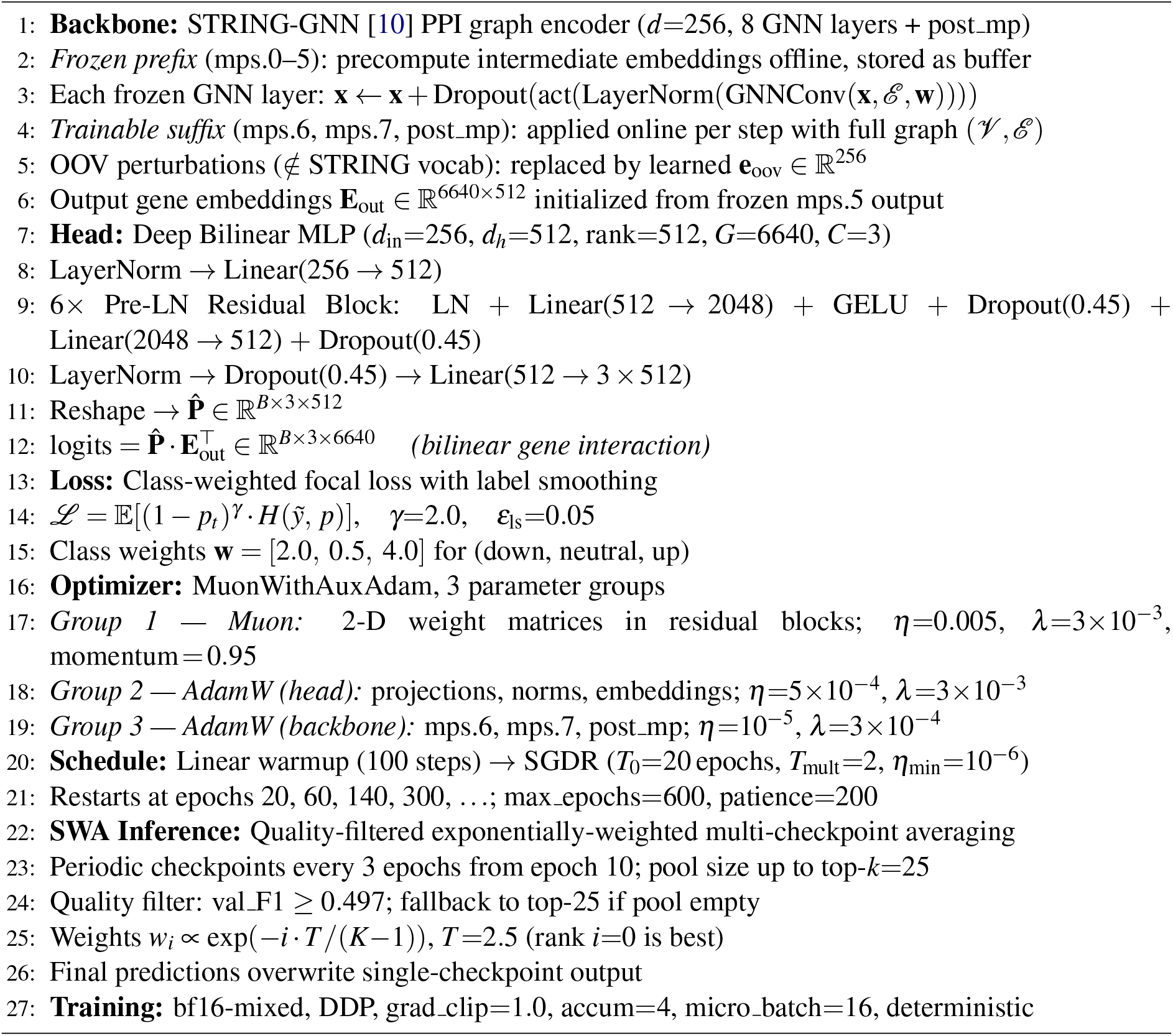

#### Synthesis and Implications

Across all four cell lines, the STRING GNN’s protein-protein interaction (PPI) representations emerged as a universal anchor for high performance [10]. STRING GNN denotes a graph neural network built over the STRING protein–protein interaction (PPI) network [24], so its backbone already encodes high-confidence interaction topology before fine-tuning. Yet the implementation varied from partial fine-tuning (hTERT-RPE1) to frozen feature extraction with complex gating (HepG2, K562). This suggests that while PPI topology provides a foundational “prior” for gene regulation, the specific regulatory logic is highly cell-type dependent. For instance, the success of the ESM2-3B [11] fusion in HepG2 versus the *SymbolEncoder* in Jurkat points to different underlying biological drivers: protein-structural constraints may dominate HepG2 responses, while Jurkat responses may be more effectively captured by broader functional gene families. These findings indicate that future foundation models for biology should not only be large in scale but also flexible enough to accommodate these cell-specific “delivery mechanisms” for structural and semantic information.

##### Algorithm 2

Discovered Architecture — HepG2 Perturbation DEG (Frozen ESM2-3B + STRING GNN, Extended Cosine Restarts)

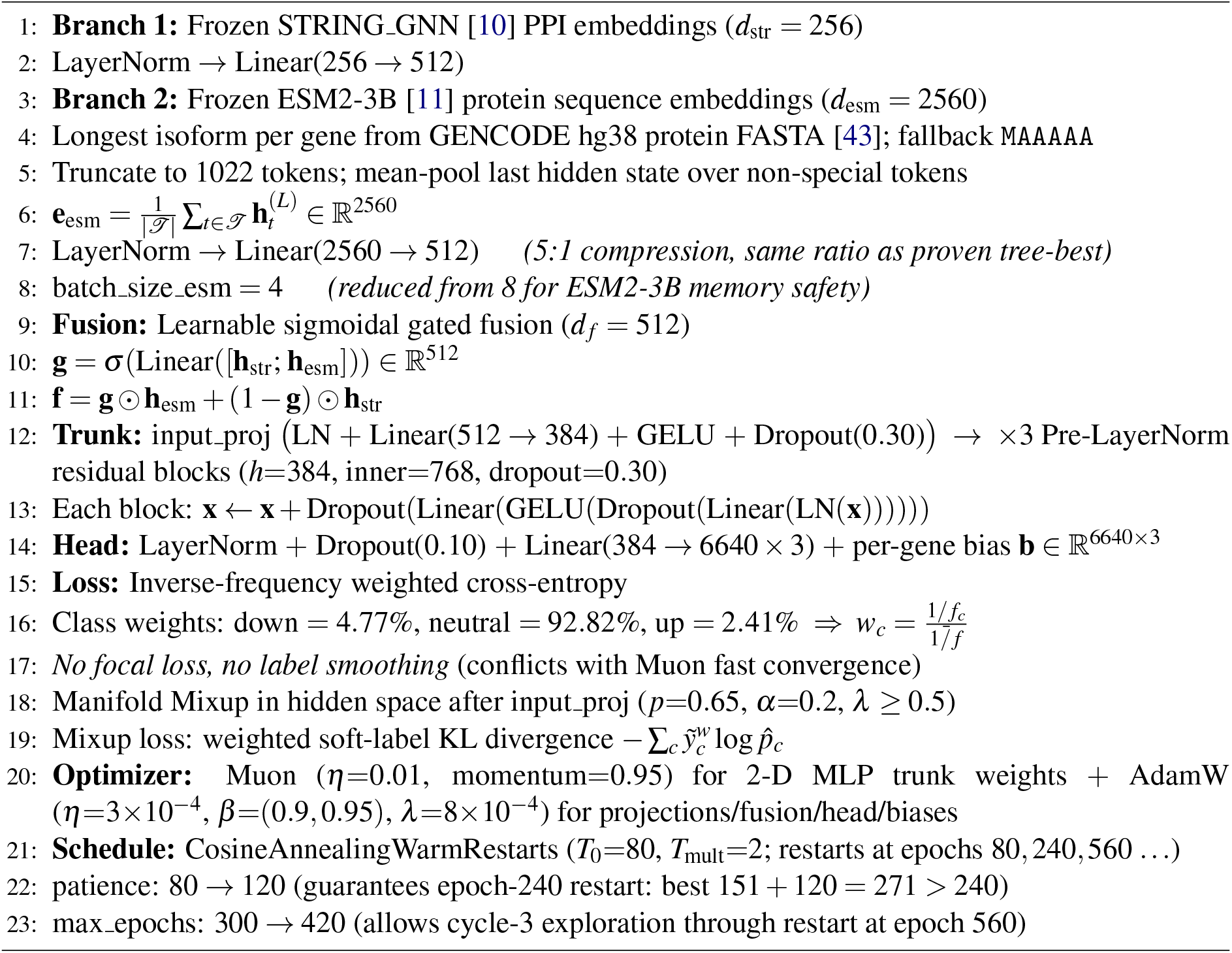

##### Algorithm 3

Discovered Architecture — Jurkat Perturbation DEG (LoRA AIDO.Cell + Frozen STRING GNN + 4-Token Cross-Attention Fusion)

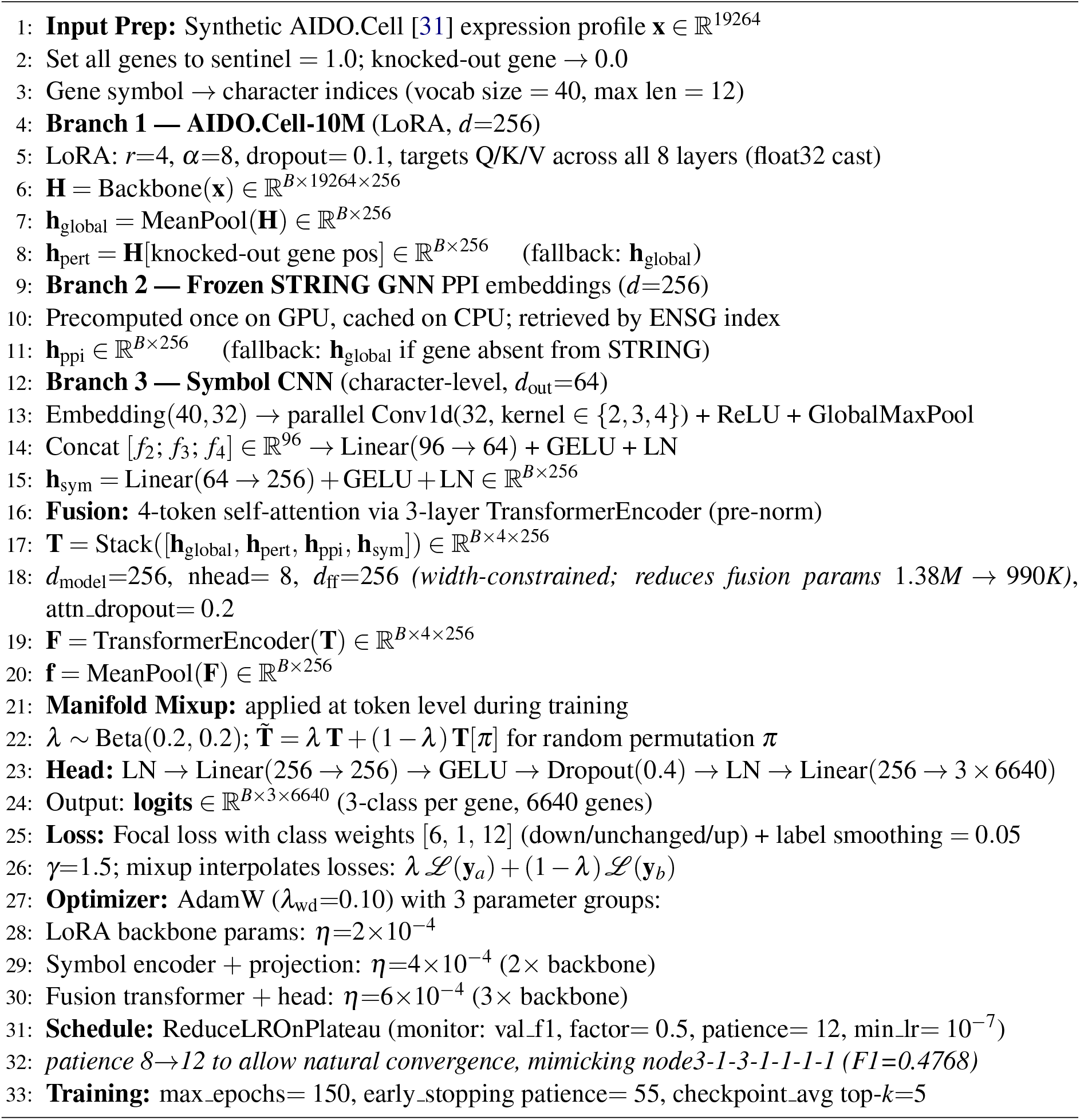

##### Algorithm 4

Discovered Architecture — K562 Perturbation DEG (AIDO.Cell-100M All-18-Layer LoRA + STRING GNN K=16 2-Head)

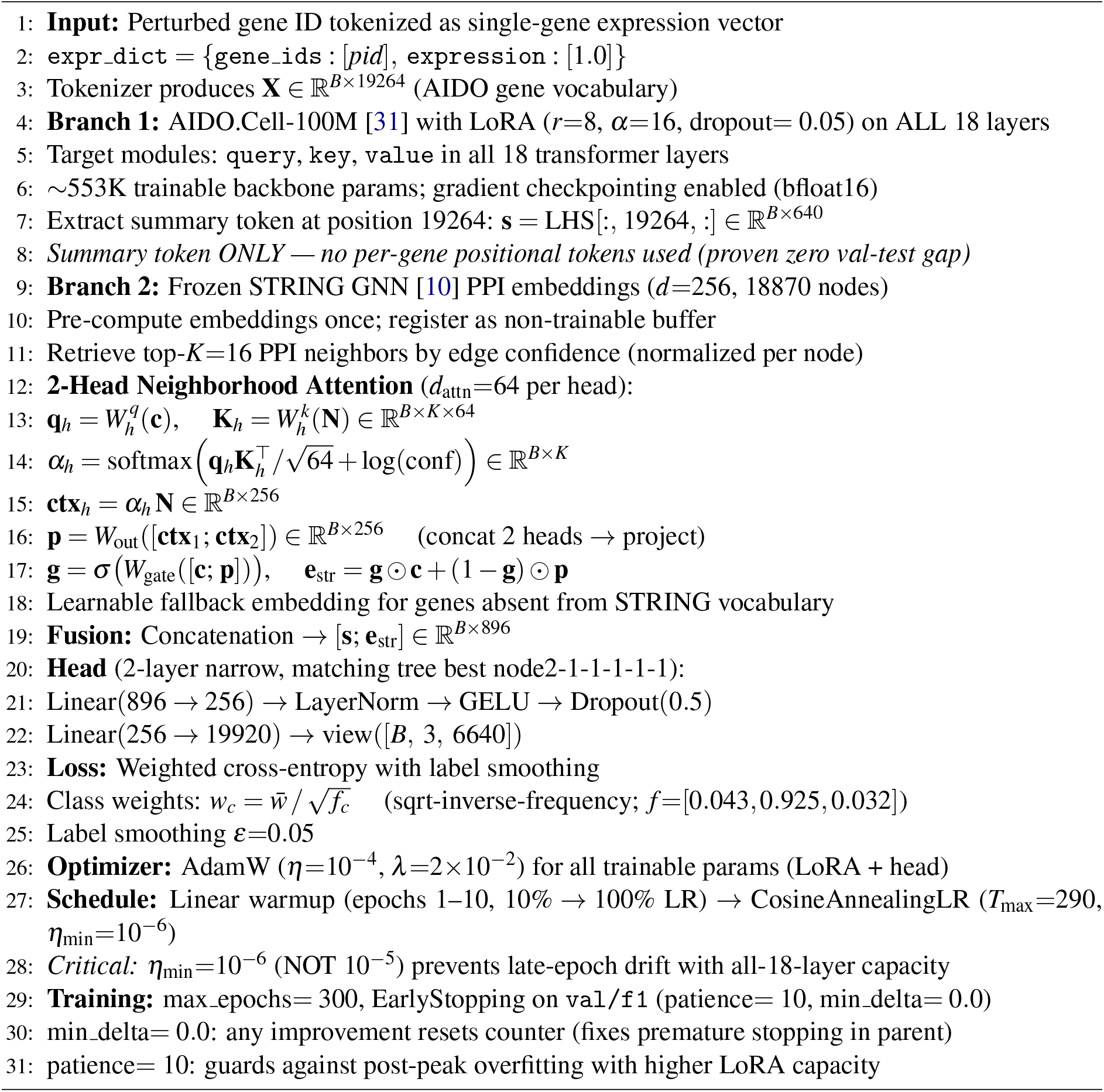

## Notes

### Competing Interest Statement

The authors have declared no competing interest.

### Summary of Updates

Fix the MPRA issues, make this paper more clear.

